# Vocal Signatures of Stress Relief: Effects of Appeasing Harness and Synthetic Pheromone on Puppy Whine Acoustics in Separation Context (*Canis familiaris*)

**DOI:** 10.64898/2026.04.02.715714

**Authors:** Romane Philippe, Anahita Le-Bourdiec-Shaffii, Vassilios Kaltsatos, David Reby, Mathilde Massenet

## Abstract

In mammals, loud, high-pitched, and harsh-sounding calls typically accompany heightened emotional arousal, particularly during distress such as separation. However, whether subtle arousal reductions can be detected through acoustic analysis within a single negative context remains unclear. We investigated whether source-related acoustic parameters of puppy whines reflect arousal modulations induced by calming interventions during maternal separation. Thirty-five eight-week-old Beagle puppies were recorded under four conditions combining synthetic appeasing pheromone and a pressure harness. Vocal behavior, activity, whine duration, and intensity, did not significantly differ across treatments, suggesting interventions did not suppress separation-related vocal responses. Nevertheless, calming products selectively altered acoustic parameters known to index arousal in dog vocalizations. Puppies receiving combined treatments produced whines with lower fundamental frequency (*f*_o_) and reduced *f*_o_ variability, while pheromone exposure increased call tonality, reflected by reduced jitter and shimmer and elevated harmonics-to-noise ratios. Spectral entropy remained unchanged, possibly because the proportion of whines containing nonlinear phenomena did not vary across conditions. Reductions in *f*_o_, *f*_o_ variability, and acoustic roughness are consistent with established correlates of lower arousal in mammals, suggesting source-related vocal parameters sensitively capture subtle arousal shifts even when overt vocal behavior remains stable, supporting their use as bioacoustic indicators for evaluating welfare interventions.

## INTRODUCTION

The objective assessment of emotional states in animals represents a central component of animal welfare sciences. Among non-invasive methods, analyzing the acoustic structure of vocalizations offers a particularly powerful approach, as vocal signals carry reliable information about the caller’s physiological and emotional condition (Briefer, 2012, 2020). Emotional states influence physiological processes (such as respiration, salivation, muscle tensions), and vocal behaviors. These physiological changes modify the configuration of the vocal apparatus and consequently alter the acoustic structure of vocalizations (Briefer, 2012, 2020).

Across mammals, elevated emotional arousal is typically associated with increased vocal activity, as reported in cows (*Bos taurus*; Thomas et al., 2001), pigs (*Sus scrofa*; Schrader & Todt, 1998; Weary et al., 1998), sheep (*Ovis aries*; Sèbe et al., 2012), cats (*Felis catus*; Yeon et al., 2011), meerkats (*Suricata suricatta*; Manser, 2001), foxes (*Vulpes vulpes*; Gogoleva et al., 2010), and marmosets (*Callithrix jacchus*; Yamaguchi et al., 2010). Beyond vocal activity, arousal also shapes the acoustic structure of calls. First, high arousal induces a shift from closed-to open-mouth vocal production, altering formant frequencies, the resonance frequencies of the vocal tract that shape the spectral envelope of the sound (sheep: Sèbe et al., 2012; cats: Shipley et al., 1991; Scheumann et al., 2012; cows: Green et al., 2020; goats (*Capra aegagrus hircus*): Ruiz-Miranda et al., 1993; Briefer et al., 2014; wapiti (*Cervus canadensis*): Reby et al., 2016; pigs: Briefer et al., 2019; Friel et al., 2019). Second, heightened states are frequently associated with increased subglottal pressure, causing the vocal folds to vibrate at higher rates and with greater amplitude, yielding vocalizations that are higher-pitched, louder, and longer (Briefer, 2012, 2020). Arousal can also affect the muscular tension applied to the larynx, altering the configuration and vibratory patterns of the vocal folds (Briefer, 2012, 2020; Muir et al., 2025). Such changes often produce irregular and harsh-sounding vocal elements known as nonlinear phenomena (NLP), including frequency jumps, subharmonics, deterministic chaos, amplitude modulation, and biphonation (Massenet et al., 2025; Anikin & Herbst, 2025) (see Figure 1 for details).

**Figure 1.**
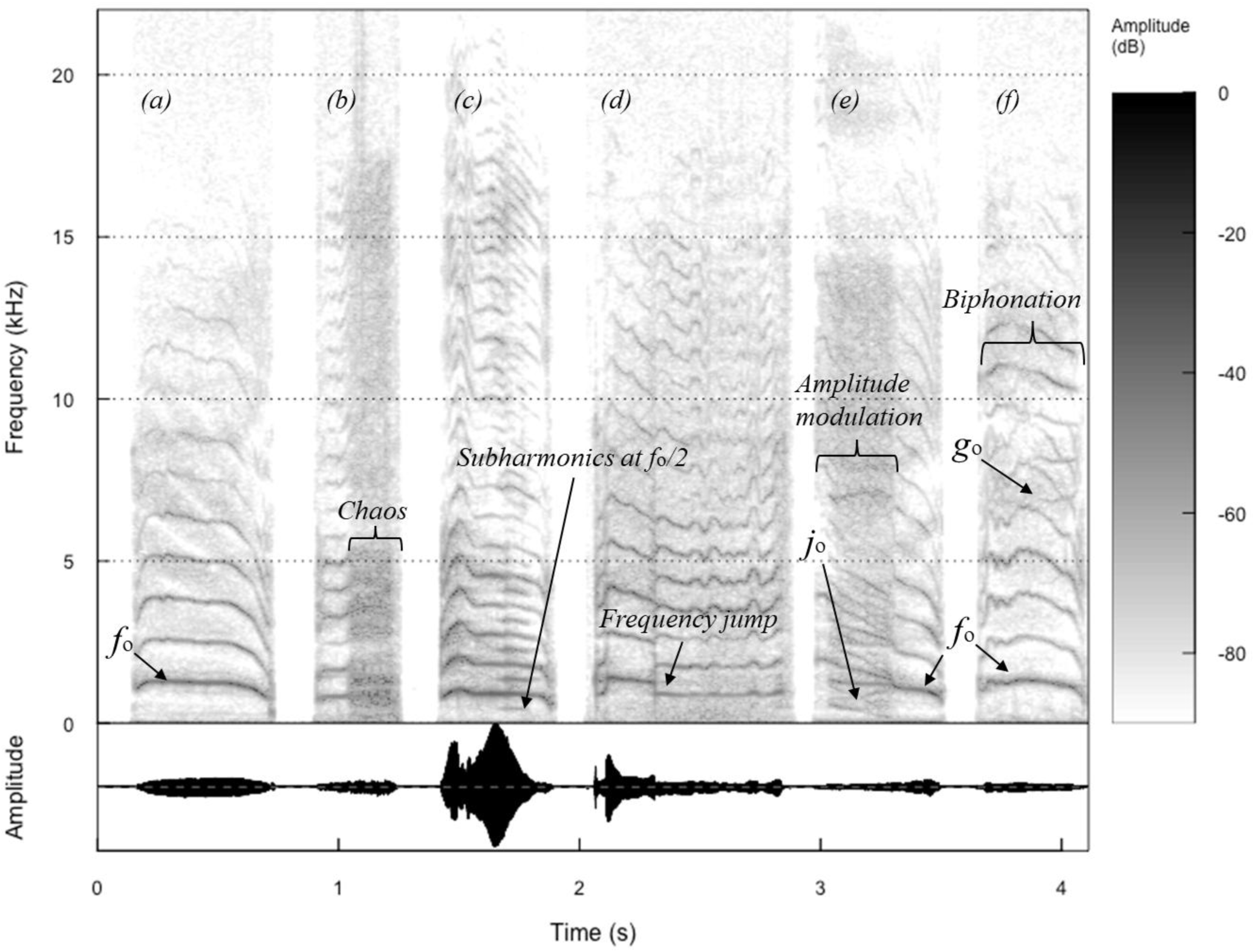
Spectrographic representations of puppy whines illustrating the different types of nonlinear phenomena (NLP). (a) A whine without NLP, showing the fundamental frequency (*f*_o_). (b) Deterministic chaos, appearing as a noisy section where *f*_o_ is difficult to track. (c) Subharmonics, corresponding to spectral bands harmonically related to *f*_o_, often occurring at *f*_o_/2. (d) Frequency jump, corresponding to an abrupt discontinuity in *f*_o_. (e) Amplitude modulation and (f) biphonation, both resulting from the vibration of a second supralaryngeal source. In amplitude modulation, spectral sidebands appear at linear combinations of m × *f*_o_ ± n × *j*_o_ (with *f*_o_ > *j*_o_), whereas in biphonation they occur at m × *g*_o_ ± n × *f*_o_ (with *g*_o_ > *f*_o_), where m and n are integers.

These emotionally driven modulations in source-related acoustic parameters (i.e., acoustic features reflecting vocal fold vibration properties, such as fundamental frequency, harmonicity, and nonlinear phenomena, as opposed to filter-related parameters corresponding to formant frequencies shaped by the vocal tract) have been documented across numerous mammalian species, providing compelling evidence that spectro-temporal features of calls function as robust markers of emotional states (Briefer 2012, 2020). Reliable correlations between acoustic parameters and stress responses have been reported in pigs (Puppe et al., 2005; Düpjan et al., 2008; von Borell et al., 2009), marmots (*Marmota flaviventris*; Blumstein, 2008), elephants (*Loxodonta africana*; Soltis, 2011), cats (Scheumann, 2012), meerkats (Townsend & Manser, 2011), cows (Green, 2020; Schnaider et al., 2021), and dogs (*Canis familiaris*; Marx, 2021; Massenet et al., 2025). For example, the roars of infant elephants produced in negatively valenced and high arousal contexts, such as a separation from their mother, have a significantly longer duration, higher fundamental frequency (*f*_o_; i.e., the rate at which the vocal folds vibrate, perceived as pitch), and higher levels of NLP than those produced during suckling, which characterizes a positively valenced and low arousal context (Soltis, 2005).

However, most past studies have focused on markedly different emotional contexts, such as contrasting high-arousal distress with positive experiences (e.g., elephants: Soltis et al., 2011; pigs: Briefer et al., 2019; Friel et al., 2019; horses: Briefer et al., 2015). These studies thus provide limited insight into whether acoustic parameters can reliably capture variations in arousal within a single emotional context. Detecting such fine-grained changes is particularly important for animal welfare research, where monitoring fluctuations in stress intensity may inform the evaluation of mitigation strategies. To address this gap, we propose to examine vocal correlates associated with reductions in distress within a stressful context, following interventions designed to regulate arousal.

Maternal separation in domestic dogs provides an appropriate model for such investigation. Brief separations from the mother and littermates usually elicit strong vocal responses in puppies, with the production of relatively high-pitched whines that function to signal urgent need for care (Bleicher, 1963; Hepper, 1994; Massenet et al., 2024). Additionally, these separation whines are often perceptually rough due to frequent occurrence of NLP (Marx et al., 2021; Massenet et al., 2022; Massenet et al., 2025). In particular, the prevalence of NLP in puppy whines increases over time following separation from the mother. This likely reflects alterations in vocal fold vibration caused by heightened laryngeal tension associated with highly aroused physiological states (Massenet et al., 2025). Moreover, greater vocal activity, characterized by increased whine duration, rate, and intensity, is directly associated with elevated arousal, as puppies vocalize more persistently in response to isolation (Pettijohn, 1977; Marx et al., 2021). Although such vocalizations are often perceived as problematic behavior, they in fact convey reliable information about the puppy’s emotional state (Massenet et al., 2025; Massenet et al., 2022), making them key for detecting variation in arousal.

To reduce stress in separation-related contexts, several calming tools have been developed, with synthetic appeasing pheromones and pressure-based devices being the most widely used in dogs. Exposures to the Dog Appeasing Pheromone (DAP), a synthetic analogue of maternal lactational secretions, have been shown to significantly decrease stress-related behaviors in numerous stressful environments or situations, such as during transportation (Gandia, 2006), veterinary interventions (Mills et al., 2006) or in shelters (Tod et al., 2005). For example, pheromone-impregnated collars reduce puppies’ activity levels (Gandia, 2006), and in veterinary clinics, pheromone diffusers limit avoidance, excessive panting, and aggression (Mills et al., 2006). Pheromone exposures are also often combined with pressure-based devices, including harnesses, which apply gentle, constant compression to the torso. The use of harnesses have shown efficacy in relieving anxiety during stressful events like thunderstorms, with owners reporting calmer behaviors characterized by an increased restlessness, and a lower vocal activity (Cottam et al., 2013; King et al., 2014). While the behavioral effects of these calming interventions are relatively well documented, their potential impacts on the acoustic structure of dog vocalizations remain unexplored. Because these tools aim to reduce arousal, they provide an ideal framework for testing whether acoustic parameters can sensitively capture within-context variation in emotional intensity.

Building on this framework, we used these products to identify bioacoustics markers of distress reduction in the whines produced by of eight-week-old Beagle puppies experiencing a short separation from their mother and littermates. We tested the independent and combined effects of synthetic appeasing pheromone and a pressure harness on source-related parameters, but not on filter-related (formants). Indeed, puppy whines are characterized by a high *f*_o,_ widely spaced harmonics and a low spectral density. As such, formants are not clearly emphasized and thus not expected to encode emotional information in this call type. In contrast, source-related parameters have been identified as significant vocal cues to arousal in puppy whines (Massenet et al., 2022, Massenet et al. 2025). Therefore, we predicted that both calming products, whether applied independently or in combination, would reduce overall whining activity and affect whine acoustics by lowering their *f*_o_ and increasing harmonicity (e.g., with less NLP).

## MATERIAL AND METHODS

### a. Subjects and ethical statements

Thirty-five Beagle puppies (15 females, 20 males) from seven litters, aged approximately eight weeks (49–56 days old), were studied at a commercial dog breeding facility in France. At this developmental stage, the puppies were weaned and were spending their final days with their mothers and littermates before adoption, which occurred at a minimum age of 58 days in accordance with French regulations. Since birth, the puppies had been housed with their mothers in dedicated enclosures, separate from other dogs.

All experimental procedures complied with French legislation on animal experimentation and welfare. The study was approved by the Ethical Committee of the ENES Lab (Approval No. E-42-218-0901) under the authority of the Direction Départementale de la Protection des Populations (Préfecture du Rhône). Protocols were specifically designed to minimize additional stress with handling limited to brief and gentle interactions. Puppies’ behavior was continuously monitored to ensure their well-being throughout the whole experiment, and were immediately reunited with their mother after each experiment. No signs of excessive distress (e.g., panting, yawning, or lip-licking; Beerda et al., 1998; Godbout et al., 2007) were observed, which otherwise would have led to the immediate interruption of the experiment.

### b. Experimental design

The independent effect of the appeasing products was evaluated through the ’*pheromone*’ treatment (exposure to Adaptil® synthetic pheromone only) and ’*harness*’ treatment (equipped with ThunderShirt® only). Combined effects were assessed in the ’*both*’ treatment (exposure to pheromone and puppy equipped with a harness). A ’*control*’ treatment where puppies were neither exposed to pheromones nor equipped with a harness served as baseline. Each puppy was tested in all four experimental conditions, presented in a randomized order over four consecutive days (one treatment per day).

To limit possible risks associated with pheromone contamination, we used two separate experimental rooms: one dedicated to treatments involving pheromone exposure (*pheromone* and *both*) and one to treatments without pheromone (*control* and *harness*). Additionally, experimenters were required to wear different laboratory coats in each room.

### c. Procedure

The experimental protocol consisted of three steps: (1) preparing experimental rooms to the assigned treatment (*Preparation* phase), (2) exposing the puppy to the treatment (*Exposure* phase), and (3) recording the puppy’s whines (*Recording* phase) (see Supplementary Methods Figure S1). Our protocol involved two experimenters, one of which blind to the tested experimental condition. The blind experimenter was not involved in the *Preparation* phase. This approach allowed us to avoid any behavioral biases toward the tested puppy during both *Exposure* and *Recording* phases. Each experimental room was identically configured, featuring the presentation of a towel, a pheromone bottle, a harness and dog crate for acoustic recordings (see below for details).

*Preparation* phase (15 min): In *pheromone* and *both* experimental conditions, eight sprays of Adaptil® were applied to one side of a towel (30 × 50 cm). We allowed a 15-minute period for complete evaporation of the product’s alcohols and diffusion of the pheromone into the environment (Adaptil, 2020). In *control* and *harness* experimental conditions, we used an unsprayed towel (serving as a placebo), and waited 15 minutes to maintain blindness of the second experimenter. At the end of the *Preparation* phase, the tested puppy came in the experimental room with the blind experimenter and was equipped with a ThunderShirt® harness (size XS) when required (*harness* and *both*).

*Exposure* phase (15 min): The tested puppy was positioned on the towel and placed on the blind experimenter’s lap, ensuring constant contact with the towel for 15 min. This duration aligned with Adaptil guidelines for pheromone exposure and allowed puppies’ habituation to the harness. The same procedure was used in *control* to ensure that the second experimenter remained fully blind to the tested condition. Puppy behavior and experimenter neutrality were monitored with video recordings (GoPro Hero 8).

*Recording* phase (10 min): At the end of the *Exposur*e phase, the puppy and towel were transferred in the dog crate (dimension: 107x70x77.5cm). Whines were audio-recorded with a Sennheiser MKH 70 directional microphone connected to a Zoom H4n digital recorder (44.1 kHz, 24-bit), positioned approximately 20 cm from the puppy’s mouth. Following each trial, the crate was disinfected with white vinegar and black soap, as recommended by pheromones manufacturers.

### d. Acoustic analysis

The dataset included 140 recordings of 10 minutes. Acoustic analyses were conducted as follows: selecting whines, measuring of acoustic parameters and performing manual annotations of NLP.

On average, whines accounted for 76% of all recorded vocalizations (N = 69,851). Other call types, such as grunts, barks, and yelps, were excluded from the analyses. The inter-call intervals were also removed to obtain recordings composed of successive whines only. This step was performed using Adobe Audition 2020 software. The number of whines produced over the 10-min recording and the cumulative duration of these whines (duration of the edited file) were extracted. Mean whine duration was then computed by dividing cumulative whine duration by the number of whines.

For each edited file, we then detected whine pitch using the *Voice Report* function of Praat software (version 6.1.16; Boersma & Weenink, 2005). The window for pitch detections was adjusted for each recording to optimize pitch tracking and ensure accurate *f*_o_ measurements. Source-related acoustic parameters were then extracted, including the fundamental frequency (mean *f*_o_ and *f*_o_ standard deviation), the vocal roughness (jitter, shimmer, and harmonics-to-noise ratio HNR), and the mean intensity (extracted using the *Get mean intensity* function) (Table 1). In addition to these acoustic measurements, the mean spectral (Wiener) entropy was extracted using an automated script based on the *analyze* function of the *Soundgen* package (Anikin, 2019) in R (version 4.0.2). Spectral entropy quantifies the degree of spectral flatness of an acoustic signal and provides a pitch-independent estimate of signal noisiness, complementing source-related measurements derived from pitch detections (e.g., HNR, jitter, shimmer).

**Table 1.** : **Description of the 15 vocal parameters measured.**

| Acoustics variables | Description |
| --- | --- |
| Number of whines | Number of whines over 10 minutes of recording |
| Mean duration of whines (sec) | Average whine duration, calculated as the total duration of whines in cleaned recordings divided by the total number of whines. |
| Mean intensity (dB) | Average sound intensity of the whines produced over 10 minutes of recording |
| Mean $f_0$ (Hz) | Mean $f_0$ of whines |
| Standard deviation (Hz) | Standard deviation of $f_0$ across whines |
| Jitter local (%) | Measurement of short-term disturbance of $f_0$, in terms of periodicity, of whines |
| Shimmer local (%) | Measurement of short-term disturbance of $f_0$, in terms of amplitude, of whines |
| Harmonics to noise ratio (%) | Harmonicity of the whine. A high HNR indicates a strong whine tonality, while a low HNR indicates a rough voice quality |
| Mean entropy | Wiener entropy, a measure of spectral flatness or disorder in the acoustic signal. Values range from 0 (pure, tonal sound with energy concentrated in narrow frequency bands) to 1 (broadband, noise-like signal with evenly distributed energy), serving as an indicator of overall tonality or aperiodicity in the whines. |
| Proportion of NLP whines (%) | Proportion of whine affected by chaos, subharmonics, frequency jumps, amplitude modulation and/or biphonation |
| Chaos proportion (%) | Proportion of whines with at least one episode of chaos |
| Subharmonics proportion (%) | Proportion of whines with at least one episode of subharmonics |
| Frequency jumps proportion (%) | Proportion of whines with at least one frequency jump |
| Amplitude modulation proportion (%) | Proportion of whines with at least one episode of amplitude modulation |
| Biphonation proportion (%) | Proportion of whines with at least one episode of biphonation |

Finally, we analyzed NLP occurrence in 20% of our full acoustic dataset. Whines were randomly selected over the 10 minutes of acoustic recording, ensuring a selection representative of puppy natural behavior where the production of NLP typically increases as emotional arousal increases (Massenet et al. 2025). Our selection resulted in manual annotations of NLP in 13,970 whines (i.e., frequency jumps, subharmonics, deterministic chaos, biphonation, and amplitude modulation, Figure 1), using the *Annotate to TextGrid* function in Praat. We then calculated the proportions of whines containing NLP (regardless of their type) and containing each type (one or more episodes of a given NLP type) (Table 1).

### e. Statistical analysis

We conducted linear mixed models (LMMs) using the *lmer* function from the *lme4* package in RStudio (version 4.0.2) to evaluate the effect of experimental conditions (or treatment) on each acoustic parameter. The models were fitted with a Gaussian distribution. They included the experimental conditions as a fixed effect, and the puppy identity and the order of experimental conditions as random effects. A full model (including both fixed and random effects) was compared to a null model (random effects only) using likelihood ratio tests (Wilks, 1938). When the overall effect of the treatment was statistically significant, post-hoc pairwise comparisons between each experimental condition were conducted using estimated marginal means (EMMs) from the *emmeans* package, with p-values adjusted via Bonferroni correction to account for multiple comparisons. Model assumptions were verified by visually inspecting the normality and homoscedasticity of the full model’s residuals. All statistical tests employed a significance threshold of α = 0.05.

## RESULTS

### Appeasement tools do not affect the overall production of whines

Our analyses revealed that the tested products did not influence the overall whining behavior of puppies. Neither the number of whines (full vs. null models: p = 0.10, Figure 2a), their duration (full vs. null models: p = 0.15, Figure 2b), nor their average intensity (full vs. null models: p = 0.57, Figure 2c) differed between the experimental conditions and the *Control* (see Supplementary Methods Table S1).

**Figure 2.**
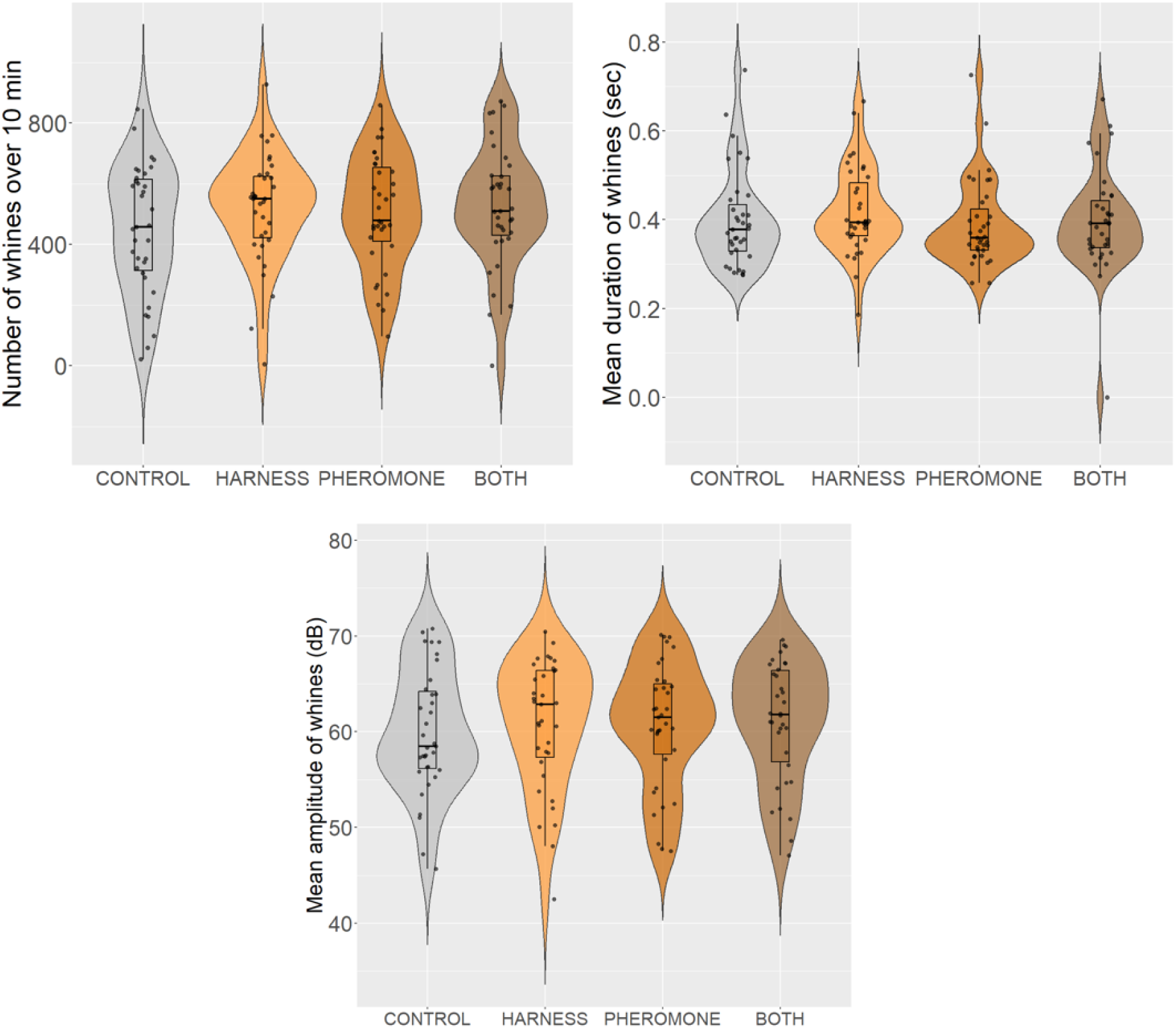
Variation in overall whine production across experimental conditions: (a) number of whines over 10 min, (b) mean whine duration, and (c) mean whine amplitude. The violin plots represent the density of data distribution for each condition, with black dots representing individual measurements. The integrated boxes with mustaches indicate the median (center line) and interquartile range (IQR, box limits), with mustaches extending up to 1.5 × IQR.

### The pressure harness reduced the mean fundamental frequency of whines and increased its stability when combined with pheromones

Given that the appeasing products did not change whine overall production, we next examined how calming products could affect finer source-related acoustic parameters of the whines such as the mean *f*_o_ and its variability. As predicted, we found that puppies produced whines at a significantly lower *f*_o_ when equipped with the harness alone (*Harness*) compared to the *Control* (estimate = 46.4 ± 15.4 Hz, p = 0.002, Table 2, Figure 3a). This effect persisted when the harness was combined with pheromone exposure (*Both*, estimate = 40.8 ± 15.7 Hz, p = 0.05). However, exposures to pheromones alone (*Pheromone*) did not affect the *f*_o_ of whines compared to the *Control* (estimate = 19.5 ± 15.4 Hz, p = 0.58). This result suggests that the harness primarily drives the reduction in *f*_o_, with a combined use with pheromones yielding a similar but slightly weaker effect. Variability in whine *f*_o_ also varied according to the tested calming product (Figure 3b, see Supplementary Methods Table S1). Puppies produced whines marked by a relatively more stable *f*_o_ (lower standard deviation) when wearing the harness and exposed to pheromones (*Both*) compared to the *Control* (estimate = 22.1 ± 8.3 Hz, p = 0.04) and to *Pheromone* (estimate = 23.1 ± 8.2 Hz, p = 0.03). However, neither the harness nor the pheromones alone affected this parameter relative to the *Control* (*Harness*, estimate = 4.5 ± 8.2 Hz, p = 0.94; *Pheromone*, estimate = -0.9 ± 8.2 Hz, p= 1), suggesting a synergistic effect when both tools are applied simultaneously. Overall, these results indicate that the harness tends to lower the mean *f*_o_, while its combination with pheromone increases *f*_o_ stability.

**Figure 3.**
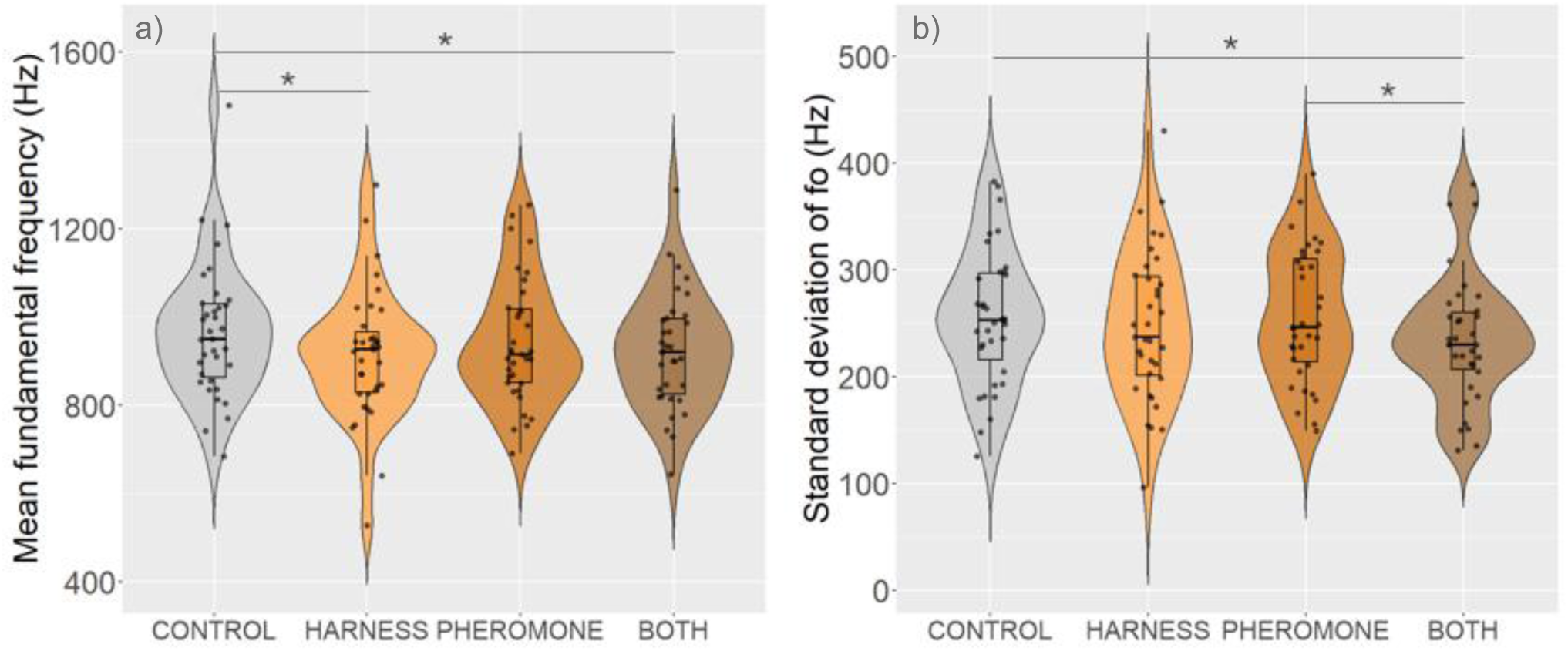
Variation in whine fundamental frequency across experimental conditions: (a) mean *f*_o_ and (b) *f*_o_ variability (standard deviation). Significant differences between conditions are indicated by p-value < 0,001 ‘***’, p-value < 0,01 ‘**’ and p-value < 0,05 ‘*’. The violin plots represent the density of data distribution for each condition, with black dots representing individual measurements. The integrated boxes with mustaches indicate the median (center line) and interquartile range (IQR, box limits), with mustaches extending up to 1.5 × IQR.

### Pheromones have mixed effects on vocal roughness

In addition to pitch modulations, we investigated parameters related to vocal roughness and spectral quality of whines (Figure 4, see Supplementary Methods Table S1). We found that puppies produced more tonal whines when exposed to or equipped with the appeasing products. Whines’ jitter was significantly lower when in *Pheromone* compared to *Control* (estimate = 0.17 ± 0.05%, p = 0.003) and in *Harness* (estimate = 0.22 ± 0.05%, p < 0.001), as well as in *Both* relative to *Harness* (estimate = 0.17 ± 0.05%, p = 0.006; Figure 4a). Furthermore, jitter did not differ significantly in *Both* compared to *Control* (estimate = 0.11 ± 0.05%, p = 0.11), confirming that pheromones, rather than the harness primarily increase vocal harmonicity.

**Figure 4.**
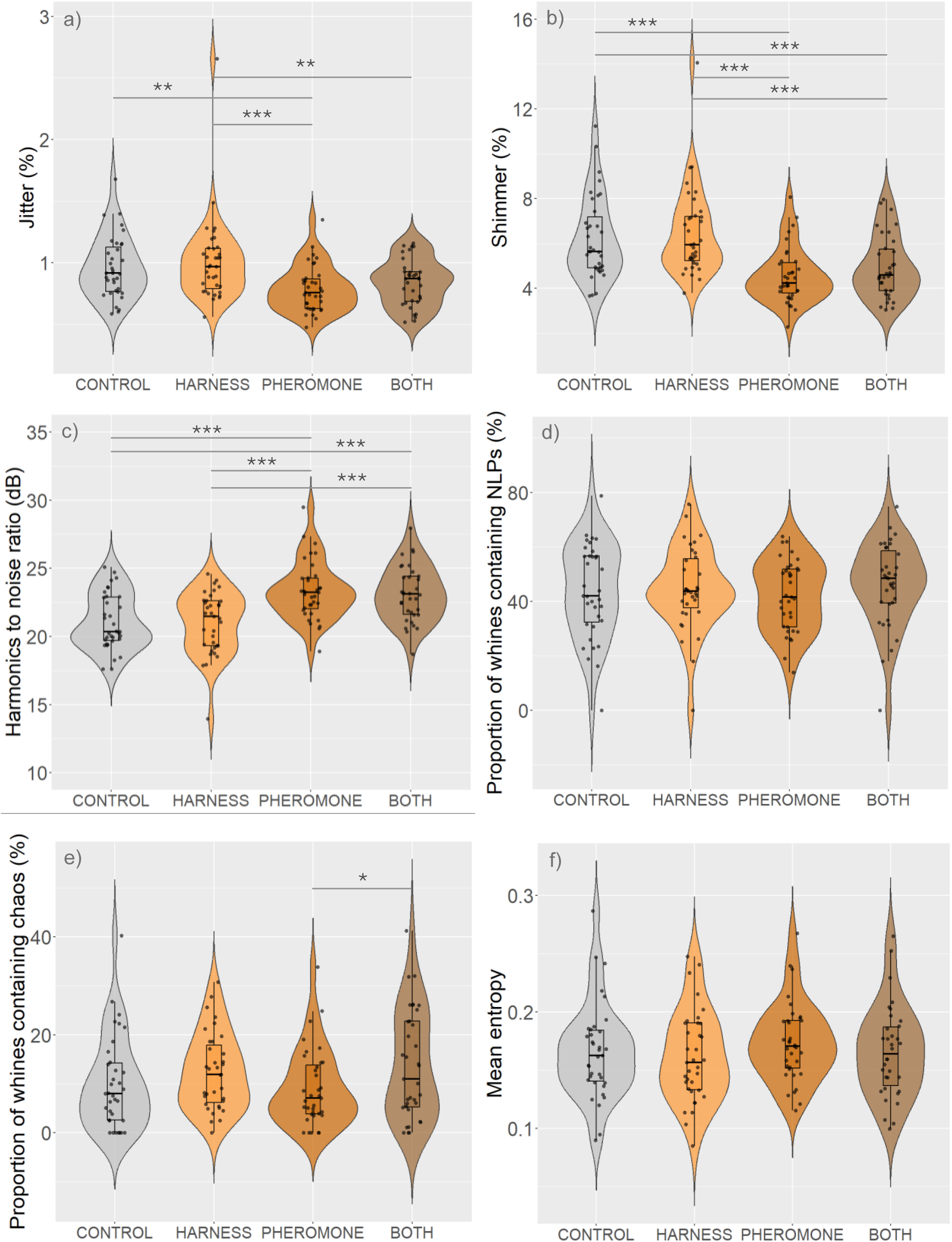
Variation in acoustic roughness of whine across experimental conditions: (a) jitter, (b) shimmer, (c) harmonics to noise ratio, (d) proportion of whine containing at least one type of NLP, (e) proportion of whines containing at least one episode of chaos and (f) mean entropy of whines. Significant differences between conditions are indicated by p-value < 0,001 ‘***’, p-value < 0,01 ‘**’ and p-value < 0,05 ‘*’. The violin plots represent the density of data distribution for each condition, with black dots representing individual measurements. The integrated boxes with mustaches indicate the median (center line) and interquartile range (IQR, box limits), with mustaches extending up to 1.5 × IQR. For all additional nonlinear parameters not presented here, see Figure S2 in the Supplementary Materials.

A similar effect was observed for the whines’ shimmer (Figure 4b): exposures to pheromones led to the production of whines characterized by a significant lower shimmer relative to those emitted in *Control* (estimate = 1.7 ± 0.3%, p < 0.001) and *Harness* (estimate = 1.9 ± 0.3%, p < 0.001). The results also show a reduction of shimmer in *Both* compared to *Control* (estimate = 1.2 ± 0.3%, p < 0.001) and *Harness* (estimate = 1.5 ± 0.31%, p < 0.001). The absence of a significant difference in shimmer between *Pheromone* and *Both* treatments (estimate = −0.4 ± 0.3%, p = 0.46), coupled with the lack of effect of the *Harness* relative to the *Control* (estimate = −0.2 ± 0.3%, p = 0.85), indicates that pheromones are responsible for the reduction of whine shimmer.

Likewise, the harmonic-to-noise ratio (HNR) (negative correlate of vocal roughness), increased significantly with treatments involving pheromones (Figure 4c). Specifically, HNR was higher in *Pheromone* compared to *Control* (*Control* – *Pheromone* estimate = -2.3 ± 0.3 dB, p < 0.001) and *Harness* (estimate = -2.3 ± 0.3 dB, p < 0.001), as well as in *Both* relative to *Control* (estimate = -2 ± 0.4 dB, p < 0.001) and *Harness* (estimate = -2.1 ± 0.4 dB, p < 0.001). We did not find significant variation in the HNR of whines emitted with the use of both two treatments involving pheromones (*Pheromone* versus *Both*; estimate = 0.3 ± 0.3 dB, p = 0.87) or between *Harness* and *Control* (estimate = 0.1 ± 0.3 dB, p = 0.98). Our results report convergent effects of pheromones across whines’ jitter, shimmer, and HNR: calls become more stable and tonal when emitted under pheromones exposures (combined or not with the harness).

Given that NLP are responsible for the perceptual roughness of vocal signals (Anikin et al., 2021), we expected a possible decrease in NLP occurrence. However, appeasing products did not significantly decrease the proportion of whines containing at least one type of NLP, regardless of the type (full vs. null models: p = 0.40; Figure 4d, see Supplementary Methods Table S1). In our subset of analyzed whines, between 42.0% and 46.2% (*Control*: 43.7 ± 17.2%; *Pheromone*: 42.1 ± 13.0%; *Harness*: 44.7 ± 15.1%; *Both*: 46.2 ± 15.5%) were affected by at least one type of NLP, indicating that NLP remain highly prevalent in puppy whines across all experimental conditions. In addition, we did not find significant variation in the proportion of whines containing specific types of NLP, except those featuring chaos. Our result show higher proportion of whines with chaos when puppies were simultaneously exposed to pheromones and equipped with the harness (*Both*) than when they were exposed to pheromones only (estimate = −4.8 ± 1.8%, p = 0.04; ; Figure 4e). However, compared to the *Control*, the proportion of whines affected by chaos did not significantly change when appeasement tools were used alone (*Pheromone* or *Harness*) or in combination (*Both*) (*Pheromone*: 0.9 ± 1.8%, p = 0.96; *Harness*: −2.4 ± 1.8%, p = 0.53; *Both*: −3.9 ± 1.8%, p = 0.15). Therefore, this result indicates a limited effect of these products on the presence of chaos in whines, which aligns with the absence of variation in the other types of NLP across treatments (see Supplementary Method Table S1, Figure S2).

Considering the absence of treatment effects on NLP occurrence, we examined the mean spectral entropy of whines, which reflects overall spectral flatness or disorder (Table 1). The treatments did not influence whine entropy (full vs. null models: p = 0.11; Figure 4f, see Supplementary Method Table S1). Across all experimental conditions, entropy remained low, ranging from 0.16 to 0.17 (*Control*: 0.16 ± 0.04; *Pheromone*: 0.17 ± 0.03; *Harness*: 0.16 ± 0.04; *Both*: 0.17 ± 0.04). This indicates that puppy whines are relatively more dominated by periodic and tonal elements than by irregular noisy ones (i.e., those involving NLP). While jitter and shimmer in whines decreased and HNR increased, the lack of variation in NLP occurrence and spectral entropy suggests that NLP episodes are likely brief and insufficiently captured by the pitch-tracking algorithms used to measure jitter, shimmer, and HNR. Consequently, appeasing products (primarily pheromones) influence pitch-based measures of vocal harmonicity but do not affect the overall spectral organization or prevalence of NLP in the whines.

## DISCUSSION

In this study, we investigated whether the effect of two commonly used putatively calming products (namely synthetic dog appeasing pheromone and a pressure harness) influence the whining behavior and whine acoustics in eight-week-old puppies experiencing a brief separation from their mother and littermates, and whether such modulations can reveal subtle variations in arousal within a single emotional context. We found that the treatments induced significant modifications in key acoustic parameters that serve as markers of emotional state in dogs, including the whine *f*_o_, its variability, and several indices of pitch-based vocal roughness (jitter, shimmer, and harmonics-to-noise ratio). However, whining rate, whine duration, amplitude, entropy, and NLP occurrence did not vary. Our results reveal a selective influence of these calming products on whine acoustics, possibly reflecting a moderate variation in the puppies’ arousal levels, and support the idea that acoustic measures can detect nuanced shifts in arousal within a same emotional context.

We first examined overall vocal activity, hypothesizing that reduced arousal would result in decreased whining rate, shorter duration of whines, or lower intensity, as evidenced in numerous mammal species (Briefer, 2012, 2020; Whitham & Miller, 2024). Contrary to our prediction, neither the pheromones nor the harness significantly affected these vocal parameters. This absence of effect aligns with previous work reporting no reduction in barking rate following pheromone exposures in shelter dogs, even if modest decreases in call amplitude were observed (Hermiston et al., 2018). Similarly, the pressure harness tested here has previously shown inconsistent effects on anxiety-related behaviors and limited impact on dog vocal behavior (Cottam et al., 2013; King et al., 2000). The persistence of whining across conditions may likely reflects the functional role of these calls, rather than an absence of product efficacy. Indeed, in young mammals, including dogs, separation calls primarily grab the mother’s attention and ultimately elicit care, making their production a robust and adaptive response to social isolation (Lingle et al 2012, Massenet et al. 2024). Taken together, our findings indicate that broad indices of vocal production reflect high arousal in vocalizers rather than gradations in its intensity. Accordingly, whining activity, as measured here, does not appear to constitute a robust parameter for evaluating the short-term effects of appeasing products in high-arousal contexts such as maternal separation.

We next examined acoustic features described as reliable indicators of arousal, such as the fundamental frequency and its variability (e.g., Briefer, 2012, 2020; Whitham & Miller, 2024). We predicted that exposures to pheromones and/or wearing the harness would reduce the whines’ mean *f*_o_ and standard deviation, given the links between elevated and unstable *f*_o_ in calls, and heightened arousal in negatively valenced contexts reported in many mammal species (Briefer, 2012, 2020). Our findings support this prediction, as puppies produced whines with a significantly lower and more stable *f*_o_ when both being exposed to pheromones and wearing the harness compared to those emitted in the control condition. Our results suggest that variation in mean *f*_o_ and standard deviation likely reflect a synergistic calming effect of products, and support the use of these parameters when evaluating arousal modulation within a given emotional context. However, while pheromones alone did not affect whine *f*_o_ and its variability, the fact that the harness alone significantly lower their *f*_o_ compared to the control points towards potential biomechanical constraints on vocal production associated with this product. The mild thoracic compression exerted by the harness may mechanically constrain respiratory and laryngeal dynamics, thereby limiting the production of higher (and more variable) frequencies (Buckley et al., 2017). Future studies combining acoustic analyses with respiratory monitoring would help disentangle emotional from biomechanical influences.

Besides *f*_o_ measures, we also examined parameters indexing vocal roughness (jitter, shimmer, HNR, NLP occurrence, and vocal entropy) given their relevance in vocal expressions of emotions in mammals (Briefer 2012, 2020). Whines produced under pheromone exposures, whether alone or in combination with the harness, exhibited lower jitter and shimmer values and higher HNR compared to the control, indicating overall higher harmonicity (and thus lower noisiness). Importantly, because the harness alone did not lead to similar changes, modulations in jitter, shimmer and HNR appear specific to the use of pheromones. Across species, increased vocal roughness has been widely reported as a correlate of high arousal or negative emotional contexts (e.g., African elephants (Soltis et al., 2011; Stoeger et al., 2011), giant pandas (*Ailuropoda melanoleuca*, Baotic et al., 2014), and domestic cats (Scheumann et al., 2012)). Within this comparative framework, the increased harmonicity observed in the whines of puppies exposed to pheromones is consistent with a reduction in arousal. Importantly, these effects occurred without changes in vocal rate, duration and intensity, indicating that pheromones primarily affect the fine structure of whines rather than overall vocal activity.

While jitter, shimmer, and HNR assess vocal roughness in solely pitch-detectable segments of vocal signals, entropy captures broader spectral disorder by measuring noisiness in both periodic and irregular sections, such as those containing NLP. We expected whine entropy and the occurrence of NLP to show patterns similar to those of jitter, shimmer and HNR. However, our results reveal that none of the tested product affected entropy and NLP occurrence. Entropy values remained relatively low (ranging from 0.16 to 0.17) and NLP occurred in between 42% and 46% of the analyzed whines (*Control*: 43.7 ± 17.2%; *Pheromone*: 42.1 ± 13.0%; *Harness*: 44.7 ± 15.1%; *Both*: 46.2 ± 15.5%). This shows that, regardless of the experimental condition, whines contain more tonal, periodic components relative to irregular noisy ones within whines, and that NLP likely affect short segments within whines. Hence, while pheromones reduced roughness in pitch-detectable segments, appeasing products did not affect broader spectral metrics. We suggest that the absence of variation in these key indicators of high arousal is due to a ceiling effect associated with the maternal separation. Indeed, previous research using a comparable maternal separation paradigm showed that NLP, especially chaos, increase with the duration of separation, indicating that these features function as indicators of escalating arousal (Massenet et al., 2022; 2025). In the present study, puppies’ separation from their mother may have induced sufficiently elevated arousal levels to limit the capacity of calming tools to produce measurable reductions in whine entropy and NLP occurrence. Therefore, our results do not challenge the significance of these parameters in evaluating animals’ emotional state, but rather, refine their interpretative scope. It indicates that both nonlinear and entropy-based metrics are particularly useful for tracking the intensification of arousal in a given context, but that they may be less sensitive to subtle downward variations once arousal has reached high levels.

### Limitations

Several limitations should be considered when interpreting the present findings. First, the sample consisted of puppies from a single breeding facility, which restricts genetic diversity and limits the generalization of our results. Future studies should include larger and more genetically diverse samples to better account for inter-individual variability in sensitivity to pheromones (Frank et al., 2010). In addition, incorporating multiple breeds would allow to determine whether the effects observed in Beagles extend more broadly across *Canis familiaris*.

Second, each puppy was exposed to each condition only once. Single, brief exposures may be insufficient for pheromone-based interventions to exert their full effects. Repeated or prolonged exposure, closer to real-world scenarios, could reveal cumulative or delayed effects that are not detected in the present design. Third, although puppies were given 15 minutes to habituate to the harness, this remained their first exposure to such equipment. Novelty or mild constraint associated with the harness may have induced additional stress rather than alleviation. Incorporating a prior familiarization phase before testing would help disentangle effects attributable to calming mechanisms from those related to novelty or physical constraint. The experimental context itself may also have influenced the results. Separation in a controlled, unfamiliar setting may have limited the sensitivity of some acoustic measures to detect subtle calming effects that could occur in more naturalistic situations. Replicating this work in ecologically relevant contexts, such as the puppy’s new home, would increase ecological validity and clarify whether similar acoustic modulations are observed under conditions that better reflect everyday experiences. Finally, methodological constraints related to acoustic measurements should be acknowledged. Jitter, shimmer, and HNR were calculated only on segments where *f*_o_ could be reliably detected. Segments containing strong nonlinearities were excluded from these analyses, which may have biased estimates of vocal roughness. Moreover, extending NLP analyses to the entire dataset would help address this limitation. Given the large number of vocalizations (>69,000), such an approach will likely require the development of automated and reliable algorithms for NLP detection and classification.

## Conclusion

This study demonstrates that exposure to calming products selectively modulate specific acoustic features of puppy whines during separation. Reductions in fundamental frequency, decreased variability, and changes in vocal roughness parameters (jitter, shimmer, and harmonics-to-noise ratio) correspond to established bioacoustic markers of lowered arousal across mammalian species. Our findings validate these source-related features as sensitive indicators of fine-grained emotional variation within a negative context. The absence of effects on overall vocal activity, spectral entropy, and the prevalence of nonlinear phenomena points out to a potential dissociation between broad behavioral measures and more subtle acoustic structure, supporting the idea that bioacoustic analyses can capture nuanced affective modulation that may not be detectable through global vocal output measures alone. By revealing this distinction, our results highlight the value of acoustic analysis as a sensitive approach for characterizing emotional expression and assessing the efficacy of calming interventions. Beyond evaluating these specific tools, the findings support bioacoustic methods as a powerful framework for identifying emotional cues in animal vocalizations. When integrated with behavioral observations, such approaches provide objective, and potentially real-time, insights into stress and welfare states without physical intervention or disruption, offering a promising avenue for improving the wellbeing of young animals.

## Supporting information

Electronic Supplementary Material

## Acknowledgements

We thank Elsa Pietrzak, Camille Toufaili, and Marine Brunel for their invaluable assistance as blind experimenters. We are also grateful to our partner breeding facility, Le Clos du Bonheur, for welcoming us and allowing the recording of their puppies. We additionally thank Floriane Fournier for creating the illustration in Figure S1.

## Data availability statement

The dataset and code are openly available at: https://zenodo.org/records/19050512?token=eyJhbGciOiJIUzUxMiJ9.eyJpZCI6Ijc0NGZkNGUwLTIzYmQtNGQ1Mi04ZDFlLWI0OTEzZDhjMTJmYyIsImRhdGEiOnt9LCJyYW5kb20iOiIxYjI1ZDY5YmEyZDBlMThhODc2OTU1MGRmMTQyNzQyZiJ9.lHSIxv85EPaz1bJDmEFxZLWiN_WflemrTzI2iJzB7BXGOnjF5B4Jnw9mvy8P_DRZaigTFwICMmgTLAbzG5orFQ. Audio recordings used in this study are not publicly available but can be provided upon reasonable request to the corresponding author.

## References

Adaptil. (2020). *Adaptil*. http://www.adaptil.com/uk

Anikin, A., & Herbst, C. T. (2025). How to analyse and manipulate nonlinear phenomena in voice recordings. Philosophical Transactions of the Royal Society B: Biological Sciences, 380(1923), Article 20240003. 10.1098/rstb.2024.0003

Anikin, A., Pisanski, K., Massenet, M., & Reby, D. (2021). Harsh is large: Nonlinear vocal phenomena lower voice pitch and exaggerate body size. Proceedings of the Royal Society B: Biological Sciences, 288(1954), Article 20210872. 10.1098/rspb.2021.0872

Baotic, A., Stoeger, A. S., Li, D., Tang, C., & Charlton, B. D. (2014). The vocal repertoire of infant giant pandas (*Ailuropoda melanoleuca*). Bioacoustics, 23(1), 15–28. 10.1080/09524622.2013.798744

Beerda, B., Schilder, M. B., van Hooff, J. A., de Vries, H. W., & Mol, J. A. (1998). Behavioural, saliva cortisol and heart rate responses to different types of stimuli in dogs. Applied Animal Behaviour Science, 58(3–4), 365–381. 10.1016/s0168-1591(97)00145-7

Bleicher, N. (1963). Physical and behavioral analysis of dog vocalizations. American Journal of Veterinary Research, 24(3), 415–426.

Blumstein, D. T., & Arnold, W. (1995). Situational specificity in Alpine-marmot alarm communication. Ethology, 100(1), 1–13. 10.1111/j.1439-0310.1995.tb00314.x

Blumstein, D. T., Richardson, D. T., Cooley, L., Winternitz, J., & Daniel, J. C. (2008). The structure, meaning and function of yellow-bellied marmot pup screams. Animal Behaviour, 76(3), 1055–1064. 10.1016/j.anbehav.2008.06.002

Boersma, P., & Weenink, D. (2005). Praat: Doing phonetics by computer (Version 5.1) [Computer software]. https://www.fon.hum.uva.nl/praat/

Briefer, E. F. (2012). Vocal expression of emotions in mammals: Mechanisms of production and evidence. Journal of Zoology, 288(1), 1–20. 10.1111/j.1469-7998.2012.00920.x

Briefer, E. F. (2020). Coding for ‘dynamic’ information: Vocal expression of emotional arousal and valence in non-human animals. In T. Aubin & N. Mathevon (Eds.), Coding strategies in vertebrate acoustic communication (Animal signals and communication, Vol. 7, pp. 137–161). Springer, Cham. 10.1007/978-3-030-39200-0_6

Briefer, E. F., Tettamanti, F., & McElligott, A. G. (2015). Emotions in goats: Mapping physiological, behavioural and vocal profiles. Animal Behaviour, 99, 131–143. 10.1016/j.anbehav.2014.11.002 (Note: This appears to be the intended reference for the cited "Briefer et al., 2014" on goats, based on verified sources; confirm if adjustment needed.)

Briefer, E. F., Vizier, E., Gygax, L., & Hillmann, E. (2019). Expression of emotional valence in pig closed-mouth grunts: Involvement of both source- and filter-related parameters. The Journal of the Acoustical Society of America, 145(5), 2895–2908. 10.1121/1.5100612

Briefer, E., Maigrot, A. L., Mandel, R., Freymond, S. B., Bachmann, I., & Hillmann, E. (2015). Segregation of information about emotional arousal and valence in horse whinnies. Scientific Reports, 5, Article 9989. 10.1038/srep09989

Buckley, L. A., & Arrandale, V. H. (2017). The use of wraps and coats for management of canine anxiety: A survey of use and perceived outcomes. Journal of Veterinary Behavior, 22, 1–7. 10.1016/j.jveb.2017.09.003

Cottam, N., Dodman, N. H., & Ha, J. C. (2013). The effectiveness of the Anxiety Wrap in the treatment of canine thunderstorm phobia: An open-label trial. Journal of Veterinary Behavior, 8(3), 154–161. 10.1016/j.jveb.2012.09.001

Düpjan, S., Schön, P. C., Puppe, B., Tuchscherer, A., & Manteuffel, G. (2008). Differential vocal responses to physical and mental stressors in domestic pigs (*Sus scrofa*). Applied Animal Behaviour Science, 114(1–2), 105–115. 10.1016/j.applanim.2007.12.005

Frank, D., Beauchamp, G., & Palestrini, C. (2010). Systematic review of the use of pheromones for treatment of undesirable behavior in cats and dogs. Journal of the American Veterinary Medical Association, 236(12), 1308–1316. 10.2460/javma.236.12.1308

Friel, M., Kunc, H. P., Griffin, K., Asher, L., & Collins, L. M. (2019). Positive and negative contexts predict duration of pig vocalisations. Scientific Reports, 9(1), Article 2062. 10.1038/s41598-019-38514-w

Gandia Estellés, M., & Mills, D. S. (2006). Signs of travel-related problems in dogs and their response to treatment with dog appeasing pheromone. Veterinary Record, 159(5), 143–148. 10.1136/vr.159.5.143

Godbout, M., Palestrini, C., Beauchamp, G., & Frank, D. (2007). Puppy behavior at the veterinary clinic: A pilot study. Journal of Veterinary Behavior, 2(4), 126–135. 10.1016/j.jveb.2007.06.002

Gogoleva, S. S., Volodina, E. V., Volodin, I. A., Kharlamova, A. V., & Trut, L. N. (2010). The gradual vocal responses to human-provoked discomfort in farmed silver foxes. Acta Ethologica, 13(2), 75–85. 10.1007/s10211-010-0076-3

Green, A. C., Clark, C. E. F., Lomax, S., Favaro, L., & Reby, D. (2020a). Context-related variation in the peripartum vocalisations and phonatory behaviours of Holstein-Friesian dairy cows. Applied Animal Behaviour Science, 231, Article 105089. 10.1016/j.applanim.2020.105089

Hepper, P. G. (1994). Long-term retention of kinship recognition established during infancy in the domestic dog. Behavioural Processes, 33(1–2), 3–14. 10.1016/0376-6357(94)90056-6

Hermiston, C., Montrose, V. T., & Taylor, S. (2018). The effects of dog-appeasing pheromone spray upon canine vocalizations and stress-related behaviors in a rescue shelter. Journal of Veterinary Behavior, 26, 11–16. 10.1016/j.jveb.2018.03.013

King, J. N., Simpson, B. S., Overall, K. L., Appleby, D., Pageat, P., Ross, C., Chaurand, J. P., Heath, S., Beata, C., Weiss, A. B., Muller, G., Paris, T., Bataille, B. G., Parker, J., & Petit, S. (2000). Treatment of separation anxiety in dogs with clomipramine: Results from a prospective, randomized, double-blinded, placebo-controlled, parallel-group, multicenter clinical trial. Applied Animal Behaviour Science, 67(4), 255–275. 10.1016/s0168-1591(00)00080-0

King, C., Buffington, L., Smith, T. J., & Grandin, T. (2014). The effect of a pressure wrap (ThunderShirt®) on heart rate and behavior in canines diagnosed with anxiety disorder. Journal of Veterinary Behavior, 9(5), 215–221. 10.1016/j.jveb.2014.06.007

Lingle, S., Wyman, M. T., Kotrba, R., Teichroeb, L. J., & Romanow, C. A. (2012). What makes a cry a cry? A review of infant distress vocalizations. Current Zoology, 58(5), 698–726. 10.1093/czoolo/58.5.698

Manser, M. B. (2001). The acoustic structure of suricates’ alarm calls varies with predator type and the level of response urgency. Proceedings of the Royal Society B: Biological Sciences, 268(1483), 2315–2324. 10.1098/rspb.2001.1773

Marx, A., Lenkei, R., Pérez Fraga, P., Bakos, V., Kubinyi, E., & Faragó, T. (2021). Occurrences of non-linear phenomena and vocal harshness in dog whines as indicators of stress and ageing. Scientific Reports, 11(1), Article 4460. 10.1038/s41598-021-83990-9

Massenet, M., Anikin, A., Pisanski, K., Reynaud, K., Mathevon, N., & Reby, D. (2022). Nonlinear vocal phenomena affect human perceptions of distress, size and dominance in puppy whines. Proceedings of the Royal Society B: Biological Sciences, 289(1973), Article 20220429. 10.1098/rspb.2022.0429

Massenet, M., Anikin, A., Pisanski, K., Reynaud, K., Mathevon, N., & Reby, D. (2024). Puppy whines mediate maternal behavior in domestic dogs. Proceedings of the National Academy of Sciences, 121(20), Article e2316818121. 10.1073/pnas.2316818121

Massenet, M., Anikin, A., Reynaud, K., Mathevon, N., & Reby, D. (2025a). Acoustic context and dynamics of nonlinear phenomena in puppy whines. In Proceedings of the International Conference on Animal Vocal Communication (preprint available at cogsci.se).

Massenet, M., Mathevon, N., Anikin, A., Briefer, E. F., Fitch, W. T., & Reby, D. (2025b). Nonlinear phenomena in vertebrate vocalizations: Mechanisms and communicative functions. Philosophical Transactions of the Royal Society B: Biological Sciences, 380(1923), Article 20240002. 10.1098/rstb.2024.0002

Mills, D. S., Ramos, D., Estelles, M. G., & Hargrave, C. (2006). A triple blind placebo-controlled investigation into the assessment of the effect of Dog Appeasing Pheromone (DAP) on anxiety related behaviour of problem dogs in the veterinary clinic. Applied Animal Behaviour Science, 98(1–2), 114–126. 10.1016/j.applanim.2005.08.012

Muir, C., Favaro, L., Massenet, M., & Reby, D. (2025). Nonlinear phenomena in mammalian vocal communication: An introduction and scoping review. Philosophical Transactions of the Royal Society B: Biological Sciences, 380(1923), Article 20240001. 10.1098/rstb.2024.0001

Pettijohn, T. F., Wong, T. W., Ebert, P. D., & Scott, J. P. (1977). Alleviation of separation distress in 3 breeds of young dogs. Developmental Psychobiology, 10(4), 373–381. 10.1002/dev.420100413

Puppe, B., Schön, P. C., Tuchscherer, A., & Manteuffel, G. (2005). Castration-induced vocalisation in domestic piglets, *Sus scrofa*: Complex and specific alterations of the vocal quality. Applied Animal Behaviour Science, 95(1–2), 67–78. 10.1016/j.applanim.2005.05.001

Reby, D., Wyman, M. T., Frey, R., Passilongo, D., Gilbert, J., Locatelli, Y., & Charlton, B. D. (2016). Evidence of biphonation and source–filter interactions in the bugles of male North American wapiti (Cervus canadensis). Journal of Experimental Biology, 219(8), 1224–1236. 10.1242/jeb.131219

Ruiz-Miranda, C. R., Szymanski, M. D., & Ingals, J. W. (1993). Physical characteristics of the vocalizations of domestic goat does *Capra aegagrus hircus* in response to their offspring’s cries. Bioacoustics, 5(1–2), 99–116. 10.1080/09524622.1993.9753232

Scheumann, M., Roser, A.-E., Konerding, W., Bleich, E., Hedrich, H.-J., & Zimmermann, E. (2012). Vocal correlates of sender-identity and arousal in the isolation calls of domestic kitten (*Felis silvestris catus*). Frontiers in Zoology, 9, Article 36. 10.1186/1742-9994-9-36

Schnaider, M. A., Heidemann, M. S., Silva, A. H. P., Taconeli, C. A., & Molento, C. F. M. (2021). Vocalization and other behaviors as indicators of emotional valence: The case of cow-calf separation and reunion in beef cattle. Journal of Veterinary Behavior, 49, 28–35. 10.1016/j.jveb.2021.11.011

Schrader, L., & Todt, D. (1998). Vocal quality is correlated with levels of stress hormones in domestic pigs. Ethology, 104(10), 859–876. 10.1111/j.1439-0310.1998.tb00036.x

Sèbe, F., Poindron, P., Andanson, S., Chandeze, H., Delval, E., Despres, G., & Boissy, A. (2012). Bio-acoustic analyses to assess emotion in animals: Acoustic patterns are linked to behavioural, cardiac and hormonal responses of ewes to the separation from their lambs. Bioacoustics, 21(1), 54.

Shipley, C., Carterette, E. C., & Buchwald, J. S. (1991). The effects of articulation on the acoustical structure of feline vocalizations. The Journal of the Acoustical Society of America, 89(2), 902–909. 10.1121/1.1894652

Soltis, J., Blowers, T. E., & Savage, A. (2011). Measuring positive and negative affect in the voiced sounds of African elephants (*Loxodonta africana*). The Journal of the Acoustical Society of America, 129(2), 1059–1066. 10.1121/1.3531798

Soltis, J., Leong, K., & Savage, A. (2005). African elephant vocal communication II: Rumble variation reflects the individual identity and emotional state of callers. Animal Behaviour, 70(3), 589–599. 10.1016/j.anbehav.2004.11.016

Stoeger, A. S., Charlton, B. D., Kratochvil, H., & Fitch, W. T. (2011). Vocal cues indicate level of arousal in infant African elephant roars. The Journal of the Acoustical Society of America, 130(3), 1700–1710. 10.1121/1.3605538

Thomas, T. J., Weary, D. M., & Appleby, M. C. (2001). Newborn and 5-week-old calves vocalize in response to milk deprivation. Applied Animal Behaviour Science, 74(3), 165–173. 10.1016/s0168-1591(01)00164-2

Tod, E., Brander, D., & Waran, N. (2005). Efficacy of dog appeasing pheromone in reducing stress and fear related behaviour in shelter dogs. Applied Animal Behaviour Science, 93(3–4), 295–308. 10.1016/j.applanim.2005.01.007

Townsend, S. W., & Manser, M. B. (2011). The function of nonlinear phenomena in meerkat alarm calls. Biology Letters, 7(1), 47–49. 10.1098/rsbl.2010.0537

von Borell, E., Bünger, B., Schmidt, T., & Horn, T. (2009). Vocal-type classification as a tool to identify stress in piglets under on-farm conditions. Animal Welfare, 18(4), 407–416. 10.1017/S0962728600000816

Weary, D. M., Braithwaite, L. A., & Fraser, D. (1998). Vocal response to pain in piglets. Applied Animal Behaviour Science, 56(2–4), 161–172. 10.1016/s0168-1591(97)00092-0

Whitham, J. C., & Miller, L. J. (2024). Utilizing vocalizations to gain insight into the affective states of non-human mammals. Frontiers in Veterinary Science, 11, Article 1366933. 10.3389/fvets.2024.1366933

Wilks, S. S. (1938). The large-sample distribution of the likelihood ratio for testing composite hypotheses. The Annals of Mathematical Statistics, 9(1), 60–62. 10.1214/aoms/1177732360

Yamaguchi, C., Izumi, A., & Nakamura, K. (2010). Time course of vocal modulation during isolation in common marmosets (*Callithrix jacchus*). American Journal of Primatology, 72(8), 681–688. 10.1002/ajp.20824

Yeon, S. C., Kim, Y. K., Park, S. J., Lee, S. S., Lee, S. Y., Suh, E. H., Houpt, K. A., Chang, H. H., Lee, H. C., Yang, B. G., & Lee, H. J. (2011). Differences between vocalization evoked by social stimuli in feral cats and house cats. Behavioural Processes, 87(2), 183–189. 10.1016/j.beproc.2011.03.003

