## Supplementary material for "Vocal Signatures of Stress Relief: Effects of Appeasing Harness and Synthetic Pheromone on Puppy Whine Acoustics in Separation Context (*Canis familiaris*)": Electronic Supplementary Material

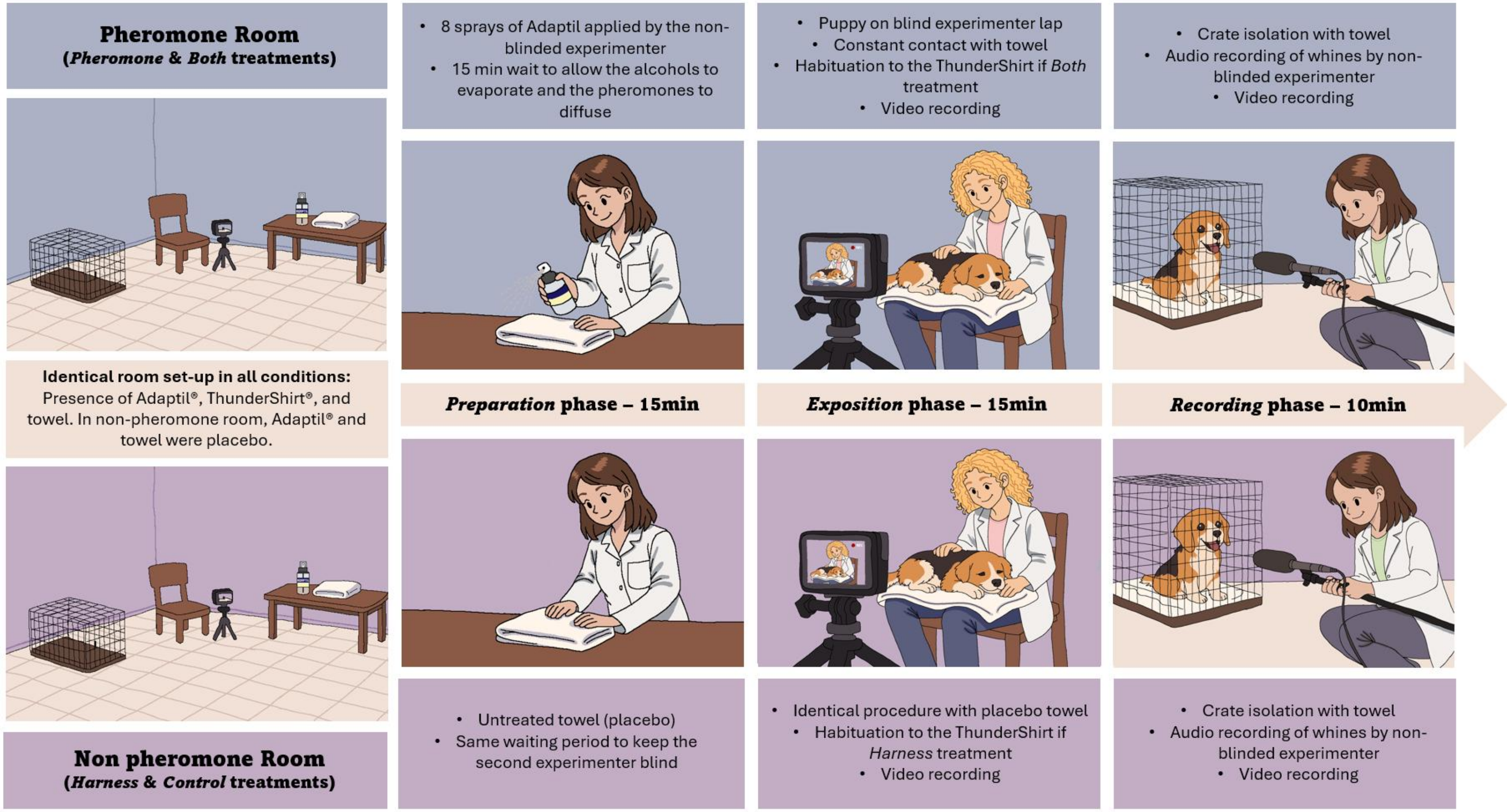


**Figure S1: Illustration of the Three-Phase Experimental Protocol (*Preparation*, *Exposure*, *Recording*)**

| Acoustic variables | Model significance | *Control* versus *Harness* | *Control* versus *Pheromone* | *Control* versus *Both* | *Harness* versus *Pheromone* | *Harness* versus *Both* | *Pheromone* versus *Both* |
|---|---|---|---|---|---|---|---|
| Number of whines | 0.10 | -54.5 ± 33.7 | -54.3 ± 33.8 | -83.5 ± 33.9 | 0.18 ± 33.6 | -29 ± 34.1 | -29.2 ± 33.6 |
| | | 0.37 | 0.38 | 0.07 | 1 | 0.83 | 0.82 |
| Average duration of whines (s) | 0.15 | -0.02 ± 0.01 | 0.01 ± 0.01 | 0 ± 0.01 | 0.03 ± 0.01 | 0.02 ± 0.01 | 0 ± 0.01 |
| | | 0.53 | 0.86 | 0.98 | 0.14 | 0.32 | 0.97 |
| Mean intensity (dB) | 0.57 | -0.8 ± 0.9 | -0.9 ± 0.9 | -1.3 ± 1 | -0.1 ± 0.9 | -0.5 ± 1 | -0.4 ± 0.9 |
| | | 0.85 | 0.78 | 0.53 | 1 | 0.94 | 0.97 |
| Mean fundamental frequency (Hz) | **0.01*** | **46.4 ± 15.4** | 19.5 ± 15.4 | **40.8 ± 15.7** | -26.9 ± 15.3 | -5.6 ± 15.8 | 21.3 ± 15.5 |
| | | **0.02*** | 0.58 | **0.05*** | 0.30 | 0.98 | 0.52 |
| Standard deviation (Hz) | **0.02*** | 4.5 ± 8.2 | -0.9 ± 8.2 | **22.2 ± 8.3** | -5.5 ± 8.1 | 17.6 ± 8.4 | **23.1 ± 8.2** |
| | | 0.94 | 1 | **0.04*** | 0.91 | 0.16 | **0.03*** |
| Jitter (%) | **< 0.001***** | -0.05 ± 0.05 | **0.17 ± 0.05** | 0.11 ± 0.05 | **0.22 ± 0.05** | **0.17 ± 0.05** | -0.06 ± 0.05 |
| | | 0.68 | **0.003**** | 0.11 | **<0.001***** | **0.006**** | 0.63 |
| Shimmer (%) | **< 0.001***** | -0.2 ± 0.3 | **1.7 ± 0.3** | **1.2 ± 0.3** | **1.9 ± 0.3** | **1.5 ± 0.31** | -0.4 ± 0.3 |
| | | 0.85 | **<0.001***** | **<0.001***** | **<0.001***** | **<0.001***** | 0.46 |
| HNR (dB) | **< 0.001***** | 0.1 ± 0.3 | **-2.3 ± 0.3** | **-2 ± 0.4** | **-2.3 ± 0.3** | **-2.1 ± 0.4** | 0.3 ± 0.3 |
| | | 0.98 | **<0.001***** | **<0.001***** | **<0.001***** | **<0.001***** | 0.87 |
| Mean entropy | 0.11 | 0.004 ± 0.004 | - 0.007 ± 0.005 | 0.002 ± 0.005 | -0.011 ± 0.005 | -0.002 ± 0.005 | 0.008 ± 0.005 |
| | | 0.81 | 0.48 | 0.98 | 0.10 | 0.96 | 0.28 |
| Proportion of whines containing NLP(%) | 0.40 | -0.9 ± 2.5 | 1.6 ± 2.5 | -2.4 ± 2.5 | 2.6 ± 2.5 | -1.5 ± 2.5 | -4.1 ± 2.5 |
| | | 0.98 | 0.91 | 0.76 | 0.72 | 0.94 | 0.35 |
| Chaos proportion (%) | **0.03*** | -2.4 ± 1.8 | 0.9 ± 1.8 | -3.9 ± 1.8 | 3.3 ± 1.8 | -1.4 ± 1.8 | **-4.8 ± 1.8** |
| | | 0.53 | 0.96 | 0.15 | 0.26 | 0.85 | **0.04*** |
| Subharmonics proportion (%) | 0.13 | -1 ± 1.2 | -1.2 ± 1.2 | -2.9 ± 1.3 | -0.1 ± 1.2 | -1.9 ± 1.3 | -1.7 ± 1.2 |
| | | 0.85 | 0.78 | 0.10 | 1 | 0.45 | 0.51 |
| Frequency jumps proportion (%) | 0.95 | 0.2 ± 1 | 0.6 ± 1 | 0.3 ± 1 | 0.4 ± 1 | 0 ± 1 | -0.3 ± 1 |
| | | 1 | 0.94 | 0.99 | 0.98 | 1 | 0.99 |
| | | -0.8 ± 1.6 | -0.2 ±1.6 | -0.8 ± 1.6 | 0.6 ± 1.6 | 0 ± 1.6 | -0.6 ± 1.6 |

| | | | | | | | |
|---|---|---|---|---|---|---|---|
| Amplitude modulation proportion (%) | 0.94 | 0.96 | 1 | 0.96 | 0.98 | 1 | 0.98 |
| Biphonation proportion (%) | 0.17 | 0.8 ± 1.3 | 2.7 ± 1.3 | 2.3 ± 1.3 | 1.9 ± 1.3 | 1.4 ± 1.4 | -0.4 ± 1.3 |
| | | 0.93 | 0.20 | 0.34 | 0.51 | 0.71 | 0.99 |

**Table S1: Results of the linear mixed models conducted on each acoustic variable and post-hoc tests realized for comparison between experimental conditions.** Estimate, standard deviation and p-value are reported. *Significant p-values are represented in bold with p-value < 0.001 '***', p-value < 0.01 '**' and p-value < 0.05 '*'*

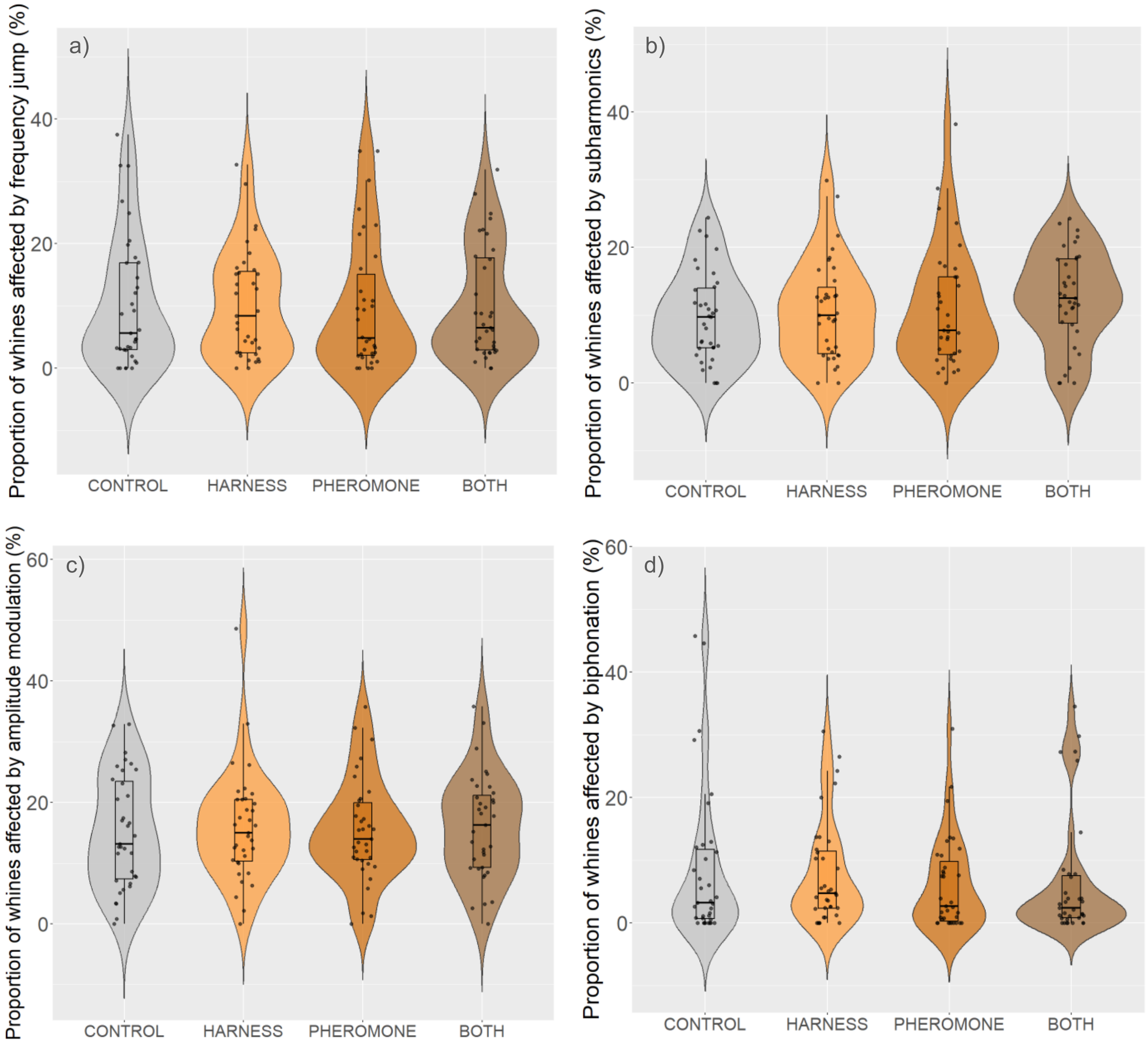


**Figure S2. Additional acoustic parameters related to the proportion of different NLP types across experimental conditions:** proportion of whines affected by (a) frequency jumps, (b) subharmonics, (c) amplitude modulation, and (d) biphonation. The violin plots represent the density of data distribution for each condition, with black dots representing individual measurements. The integrated boxes with mustaches indicate the median (center line) and interquartile range (box limits), with mustaches extending up to 1.5 × IQR.
